# Genome-wide comparison of toxigenic and non-toxigenic *Corynebacterium diphtheriae* isolates identifies differences in the pan genomes between respiratory and cutaneous strains

**DOI:** 10.1101/143800

**Authors:** Verlaine J Timms, Trang Nguyen, Taryn Crighton, Marion Yuen, Vitali Sintchenko

**Author notes:** Postal Address: Centre for Infectious Diseases and Microbiology, Westmead Hospital, PO Box 533 Wentworthville NSW 2145, Australia.

## Abstract

**Objectives:** *Corynebacterium diphtheriae* is the main etiological agent of diphtheria, a global disease causing life-threatening infections, particularly in infants and children. Vaccination with diphtheria toxoid protects against infection with potent toxin producing strains. However a growing number of apparently non-toxigenic but potentially invasive *C. diphtheriae* strains are identified in countries with low prevalence of diphtheria, raising key questions about genomic structures and population dynamics of the species.

**Methods:** This study examined genomic diversity among 47 *C. diphtheriae* isolates collected in Australia over a 10-year period using whole genome sequencing. Phylogeny was determined using SNP-based mapping and genome wide analysis.

**Results:** *C. diphtheriae* sequence type (ST) 32, a non-toxigenic ST with evidence of enhanced virulence that is also circulating in Europe, appears to be endemic in Australia. Isolates from temporospatially related patients displayed the same ST and similarity in their core genomes. The genome-wide analysis highlighted a role of pilins, adhesion factors and iron utilization in infections caused by toxigenic as well as non-toxigenic strains.

**Conclusions:** The genomic diversity of toxigenic and non-toxigenic strains of *C. diphtheriae* in Australia suggests multiple local and overseas sources of infection and colonisation. Our findings suggest that regular genomic surveillance of co-circulating toxigenic and non-toxigenic *C. diphtheriae* can deliver highly nuanced data in order to inform targeted public health actions and policy for predicting the future impact of this highly successful pathogen.

## Introduction

Diphtheria was the number one cause of infant death prior to the introduction of toxoid vaccines and since this time, infection with *Corynebacterium diphtheriae*, the causative agent of diphtheria, has been a rare challenge in the developed world (1). Toxin-negative isolates of *C. diphtheriae* have been revealed to be associated with prosthetic and native valve endocarditis and significantly, have been increasingly detected in clinical samples (2, 3). This emerging increase is thought to be due to the uptake of matrix-assisted laser desorption/ionisation time-of-flight mass spectrometry (MALDI-TOF MS) identification, but the scenario is unclear. The growing numbers of apparently non-toxigenic but potentially invasive *C. diphtheriae* isolates identified by diagnostic and public health laboratories in countries with low prevalence of diphtheria raises concerns about other virulence factors and population dynamics of the species. Since the publication of the first complete genome sequence of *C. diphtheriae* (4) the phylogeographical structure of this species and the role of iron-uptake systems, adhesions and fimbrial proteins in virulence have become key questions that need addressing (5). Furthermore, the re-emergence of potent toxigenic variants may emerge from local strains being lysogenized by a toxin gene carrying corynebacteriophage (6).

The most virulent *C. diphtheriae* are those that possess the toxin gene and produce diphtheria toxin (DT), one of the most potent exotoxins known. In order for *C. diphtheriae* to produce DT it must be lysogenized with corynebacteriophages carrying the DT gene (*tox*). The two most common bacteriophages known to infect *C. diphtheriae* are corynephage β and ω. However corynebacteriophages from *C. ulcerans* can also carry *tox* gene homologs and it remains unknown whether bacteriophages of *C. ulcerans* can lysogenize *C. diphtheriae* (7). In addition, very little is known whether other toxin gene variants exist and the role that other *C. diphtheriae* pathogenic factors may play. Toxin production is regulated by a repressor on the bacterial chromosome, DtxR in response to the amount of free iron available in the local environment (8). DtxR is also responsible for the regulation of a wide range of other genes used for colonisation, nutrient acquisition and persistence and has been shown to vary among strains (9). Such factors involved in iron metabolism and virulence determinants such as pili may also vary among strains and contribute to the success of certain clones (10).

The advancement of whole genome sequencing has led to revolutionary change for public health laboratory surveillance, opening up the potential to describe outbreaks in high resolution and to explore potential transmission routes (7, 8). The aim of this study was to examine genomic variation among *C. diphtheriae* isolates identified in the most populous state of Australia and referred to our laboratory for diphtheria toxin testing over a 10 year period. We examined whether *C. diphtheriae* transmission has been occurring locally and whether recent strains contained toxin homologs undetectable with existing assays. We also investigated if other pathogenic factors such as pili variation were contributing to possible local transmission.

## Methods

### Bacterial isolates and molecular subtyping

All *C. diphtheriae* clinical isolates collected between January 2004 and January 2016 by the public health reference laboratory at the Centre for Infectious Diseases and Microbiology, Pathology West, Westmead Hospital were included in the study. They were cultured on horse blood agar and incubated aerobically at 37°C. Bacterial isolates were identified as *C. diphtheriae* based on MALDI-TOF (cut-off 2.2) and the biotype determined biochemically using the API^®^ Coryne Strip (API bioMérieux). Toxin studies were carried out using the modified Elek test (11) and PCR for the diphtheria toxin gene (12).

### DNA extraction and whole genome sequencing (WGS)

Genomic DNA was extracted from pure cultures using the DNeasy Blood & Tissue Kit (QIAGEN). Type strains *C. diphtheriae* ATCC 27010 (C7(-)) and ATCC 13812 (PW8) were included for comparison of assembly and typing pipelines. Paired-end indexed libraries of 150bp in length were prepared from an input of 1 ng of purified DNA with the Nextera XT kit (Illumina) as per manufacturer’s instructions. DNA libraries were then sequenced using the NextSeq 500 Instrument (Illumina).

### Genome assembly and analysis

The quality of the sequence data was assessed using FastQC. Sequencing reads were assembled with Spades (13) and annotated with Prokka (14). Multiple locus sequence typing (MLST) was performed with seven loci by uploading assembled fasta sequences to the PubMLST *Corynebacterium diphtheriae* database (<u>http://pubmlst.org/cdiphtheriae/</u>). In addition, pangenome assessment and visualisation was performed using Roary (15) including alignment using MAFFT (Katoh, Misawa, Kuma, & Miyata, n.d.) and tree building with FastTree (Price et al., 2010).

To identify SNPs, fastq files were imported into Genious (8.0.4) and mapped to the reference *C. diphtheriae* NCTC 13129 using the bwa plugin (version 0.7.10). Quality based variant detection was performed using CLC Genomics Workbench v.7.0 (CLC bio Aarhus, Denmark). Variant detection thresholds were set for a minimum coverage of 10 and minimum variant frequency of 75%. SNPs were excluded if they were in regions with a minimum fold coverage of <10, within 10-bp of another SNP or <15-bp from the end of a contig. Maximum likelihood phylogenetic trees were constructed from SNP matrices using the GTR model with 100 bootstrap replications.

Antibiotic resistance was predicted using ResFinder (16). BLAST comparisons to search for toxin homologs was performed with the following toxin homologs: Corynephage beta A and B subunit (NCBI accession number P00588), Corynephage omega Diphtheria toxin (accession number P00587), Corynephage beta Diphtheria toxin homolog (accession number P00589) and *C. ulcerans* Diphtheria toxin homologs (accession number Q6YIX9 and Q5IL09). The homology of *dtx*R and all the genes from the Spa pilus clusters (A, D &H) was also determined using the BLAST suite with orthologs defined as those that had at least 50% length compared to corresponding gene in NCTC13129. The genomic data have been deposited in the NCBI Sequence Read Archive (SRA) (<u>http://www.ncbi.nlm.nih.gov/Traces/sra/</u>) under accession number (XXXX).

## Results

Forty seven isolates were included in the study. They were recovered from patients between 2004 and 2016 with 36 of these collected in 2014-2016. Three isolates were toxigenic (all biovar. *mitis*) and identified in 2015 (Table 1). The genome size was in the range of 2.2-2.7 Mb with an average G+C ratio of 53%. MLST typing revealed that some isolates shared the same or had similar sequence type (ST). Core genome analysis on *de novo* assembled genomes identified 1384 core genes, 354 ‘soft core’ genes (present between 95% - 99% of strains), 888 ‘shell genes’ (present in 15%-95% of strains), 5177 cloud genes (present between 0-15% of strains) and a total pangenome of 7803 genes. Only those strains that were known to be toxin positive (5007, 2562 and 2767B) demonstrated the toxin gene or any homologs by BLAST. No variability was observed in the *dtxR* gene for any strain in this study.

**Table 1:** *C. diphtheriae* strains analysed in this study.

| Isolate | Biotype | Year isolated | Origin | Sequence Type |
| --- | --- | --- | --- | --- |
| 3322 | *gravis* | 2007 | Cutaneous (foot) | 239 |
| 2347 | *gravis* | 2007 | Blood | 122 |
| 2235 | *mitis* | 2008 | Cutaneous (leg) | 86 |
| 3846 | *gravis* | 2009 | Respiratory | New** |
| 2682 | *gravis* | 2012 | Blood | 122 |
| 1138 | *mitis* | 2012 | Cutaneous (leg) | New** |
| 4705 | *mitis* | 2012 | Cutaneous (site unknown) | New** |
| 4447 | *gravis* | 2012 | Respiratory | 32 |
| 548 | *mitis* | 2012 | Cutaneous (site unknown) | New** |
| 2686 | *gravis* | 2013 | Respiratory | 32 |
| 3876 | *gravis* | 2013 | Cutaneous (leg) | 240 |
| 3967 | *mitis* | 2013 | Cutaneous (site unknown) | New** |
| 4089 | *gravis* | 2014 | Blood | New** |
| 4170 | *mitis* | 2014 | Cutaneous (leg) | New** |
| 3590 | *mitis* | 2014 | Cutaneous (foot) | New** |
| 2379 | *mitis* | 2014 | Cutaneous (foot) | 259 |
| 4867 | *mitis* | 2014 | Cutaneous (hand) | New** |
| 4491 | *mitis* | 2014 | Cutaneous (site unknown) | New** |
| 4650 | *mitis* | 2014 | Cutaneous (site unknown) | New** |
| 2245 | *mitis* | 2014 | Cutaneous (site unknown) | New** |
| 4218 | *gravis* | 2014 | Respiratory | 32 |
| 4526 | *gravis* | 2014 | Respiratory | New** |
| 4711 | *gravis* | 2014 | Respiratory | New** |
| 2628 | *gravis* | 2014 | Cutaneous (arm) | New** |
| 4179 | *mitis* | 2014 | Cutaneous (leg) | New** |
| 2510 | *mitis* | 2014 | Cutaneous (site unknown) | New** |
| 1425 | *mitis* | 2014 | Cutaneous (leg) | 20 |
| 2349 | *gravis* | 2014 | Cutaneous (site unknown) | 147 |
| 257 | *mitis* | 2015 | Cutaneous (site unknown) | New** |
| 2562* | *gravis* | 2015 | Cutaneous (foot) | 120 |
| 5007* | *gravis* | 2015 | Cutaneous (site unknown) | 59 |
| 4435 | *gravis* | 2015 | Cutaneous (site unknown) | New** |
| 4442 | *mitis* | 2015 | Cutaneous (site unknown) | New** |
| 4801 | *mitis* | 2015 | Cutaneous (site unknown) | New** |
| 2352 | *gravis* | 2015 | Cutaneous (leg) | New** |
| 2767A | *mitis* | 2015 | Cutaneous (ankle) | New** |
| 2767B* | *gravis* | 2015 | Cutaneous (ankle) | 381 |
| 4455 | *mitis* | 2015 | Cutaneous (site unknown) | New** |
| 4375 | *mitis* | 2015 | Cutaneous (site unknown) | New** |
| 5239 | *mitis* | 2015 | Cutaneous (penile ulcer,<br>underlying syphilis) | 6 |
| 5117 | *mitis* | 2015 | Cutaneous (site unknown) | New** |
| 3870 | *mitis* | 2015 | Cutaneous (site unknown) | 5 |
| 3888 | *gravis* | 2016 | Cutaneous (wound<br>unknown site) | 240 |
| 3067 | *mitis* | 2016 | Cutaneous | New** |
| 993 | *mitis* | 2016 | Cutaneous (foot) | New** |
| 1436 | *mitis* | 2016 | Cutaneous | New** |
| 4809 | *mitis* | 2016 | Cutaneous (penis,<br>circumcision wound) | New** |
| 1405 | *mitis* | 2016 | Cutaneous (site unknown) | New** |
| ATCC13812 | *gravis* | 1896 | Respiratory | 44 |
| ATCC27010 | *mitis* | 1954 | Respiratory | 26 |

Note: \*denotes toxin positive strains;
\*\*all new sequence types are unique (see Supplementary Table 1).

Four out of six clusters appeared to be geographically linked (Figure 1). Cluster 1 contained strains 2686, 4218, 4447 and 4526/4711 (from the same patient). This cluster had the *in silico* MLST profile for ST32; *atpA*-3, *dnaE*-1, *dnaK*-18, *fusA*-4, *leuA*-13, *odhA*-3, *rpoB*-5, with the exception of strains 4526/4711 that differed by one nucleotide (C to T) in the *dnaK* locus (Table 1 and Supplementary table 1). All isolates were from respiratory samples of adult patients residing in the same region.

**Figure 1:**
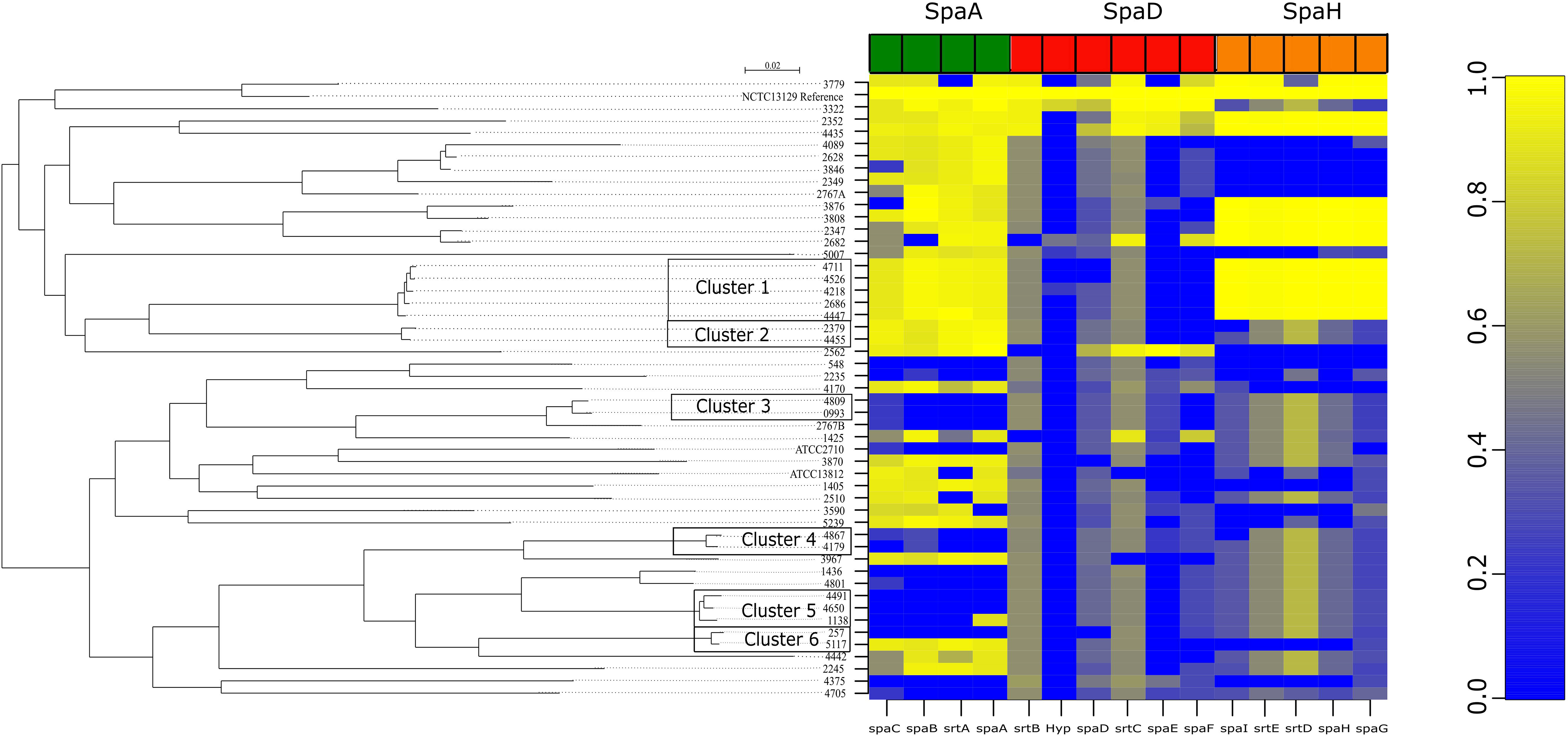
Maximum likelihood tree based on SNP detection of reads mapped to reference NCTC13129. Branch lengths correspond to numbers of nucleotide substitutions per site. The heatmap shows *spa* gene clusters when compared to the reference NCTC11329 with high homology shown in yellow, absence or poor homology shown in blue.

Analysis of the pangenome showed that cluster 1 contained unique genes denoted by I and II (Figure 2). The first (I) were mainly hypothetical proteins however two were annotated as transposable elements, one ATPase and a modification methylase that was not found in any other isolates in this study (Figure 2). An additional 11 genes (II) were also unique and mostly hypothetical but again contained transposons, unique putative outer membrane proteins and phanazine biosynthesis protein (PhzF family). Analysis of SpaA, SpaD and SpaH pilus gene clusters showed that all were present in this cluster, even though the SpaD gene cluster had low homology to the reference strain (Figure 1).

**Figure 2:**
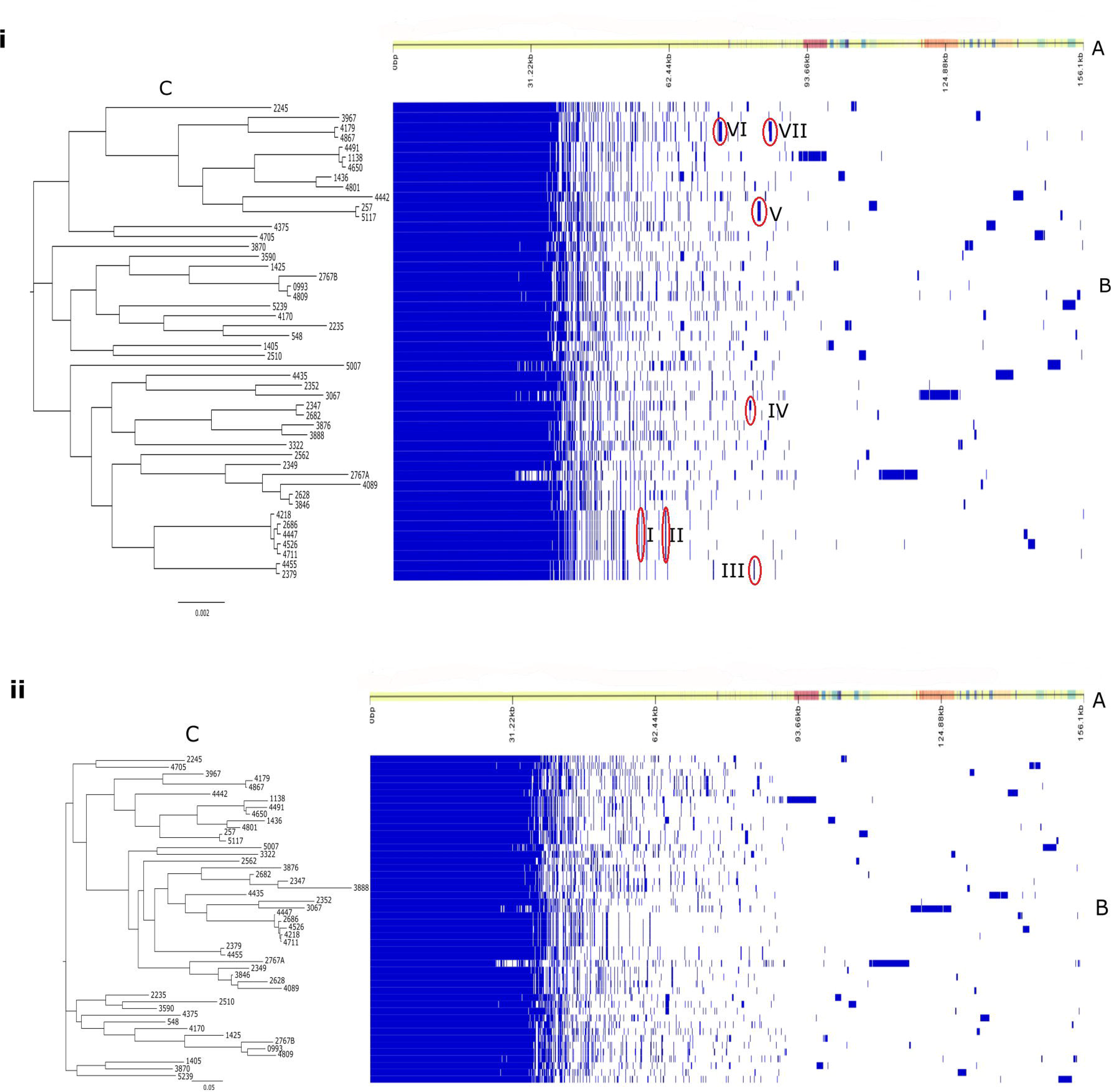
The pan genome with core genome phylogenetic tree (i) or accessory tree (ii). The top panel (A) shows the contigs inferred from the pan-genome with ordering according to pan-genome content. The middle panel (B) displays presence (blue) or absence (white) of blocks relative to genes and contigs in the pangenome. The phylogenetic tree (C) displays phylogeny based on the core (i) or accessory (ii) genome.

Cluster 2 consisted of two related isolates. The first isolate 2379 was ST259 (*atpA*-3, *dnaE*-1, *dnaK*-12, *fusA*-1, *leuA*-42, *odhA*-16, *rpoB*-31), while isolate 4455 differed by one nucleotide in the *dnaK* locus. Both isolates were predicted to be resistant to Phenicol, Sulphonamide and Tetracycline. Pangenome analysis showed that the two isolates from this cluster contained the *tetO* gene predicting resistance to tetracycline (III Figure 2). This cluster did not contain the *spaE* or *spaF* gene and had variable homology in SpaH (Figure 1).

Cluster 3 consisted of isolates 4809 and 0993 and both isolates were from teenage males. No geographic link was determined. Pangenome analysis did not show any unique genes common to both strains. Like cluster 2, strains from this cluster did not contain the *spaF* gene and had variable homology in *both* SpaD and SpaH (Figure 1).

Cluster 4 was represented by isolates 4867 and 4179 from two patients (both the same age) of the same address. The two isolates represented a new ST (*atpA*-13, *dnaE*-2, *dnaK*-32, *fusA*-33, *leuA*-no match, *odhA*-1, *rpoB*-21). These strains had two regions of genes that were unique. The first region (VI) was a phage and the second (VII) contained genes for a fimbrial subunit type 1 as well as sulphur carrying protein (ThiS), inner membrane transporter protein (RhIA) and VRR-NUC domain protein to name a few. All three Spa pilus gene clusters were highly variable in this cluster and did not have significant homology with the reference NCTC 13129 (Figure 1).

In the fifth cluster, isolates 4491, 4650 and 1138 represented a new ST with identical loci. All isolates were from males (age range 20-88 years). No markers of antibiotic resistance were observed and no unique genes were identified in this cluster. Similar to cluster 4, all three Spa pilus gene clusters were highly variable in this cluster and did not have significant homology with the reference NCTC 13129 (Figure 1).

Cluster 6 consisted of isolates 257 and 5117 and no antibiotic resistance genes were recorded. Pangenomic analysis (Figure 2, V) demonstrated unique genes for these two strains that contained *sdpA* and *sdpB*, both sporulation-delaying proteins, an integrase, von Willebrand factor type A domain protein and a putative transposon Tn552, all of which were shown to be part of a non-ribosomal peptide/ polyketide synthase cluster unique to these strains. In addition, these strains contained a large NRPS cluster with homologs to *mbtB, irtA* and *irtB*, genes known for iron regulation and survival in *M. tuberculosis* (17).

Interestingly, NRPS/PKS clusters contained small variations among strains, however most notable was an additional gene present in the Type 1 PKS cluster that was present in strains associated with systemic and cutaneous infections and was absent in respiratory isolates. This protein was annotated as a putative collagen binding protein and showed high homology (89-94%) to similar proteins in *C. diphtheriae* isolates from cutaneous infections but low homology (<84%) in respiratory strains (Figure 3).

**Figure 3.**
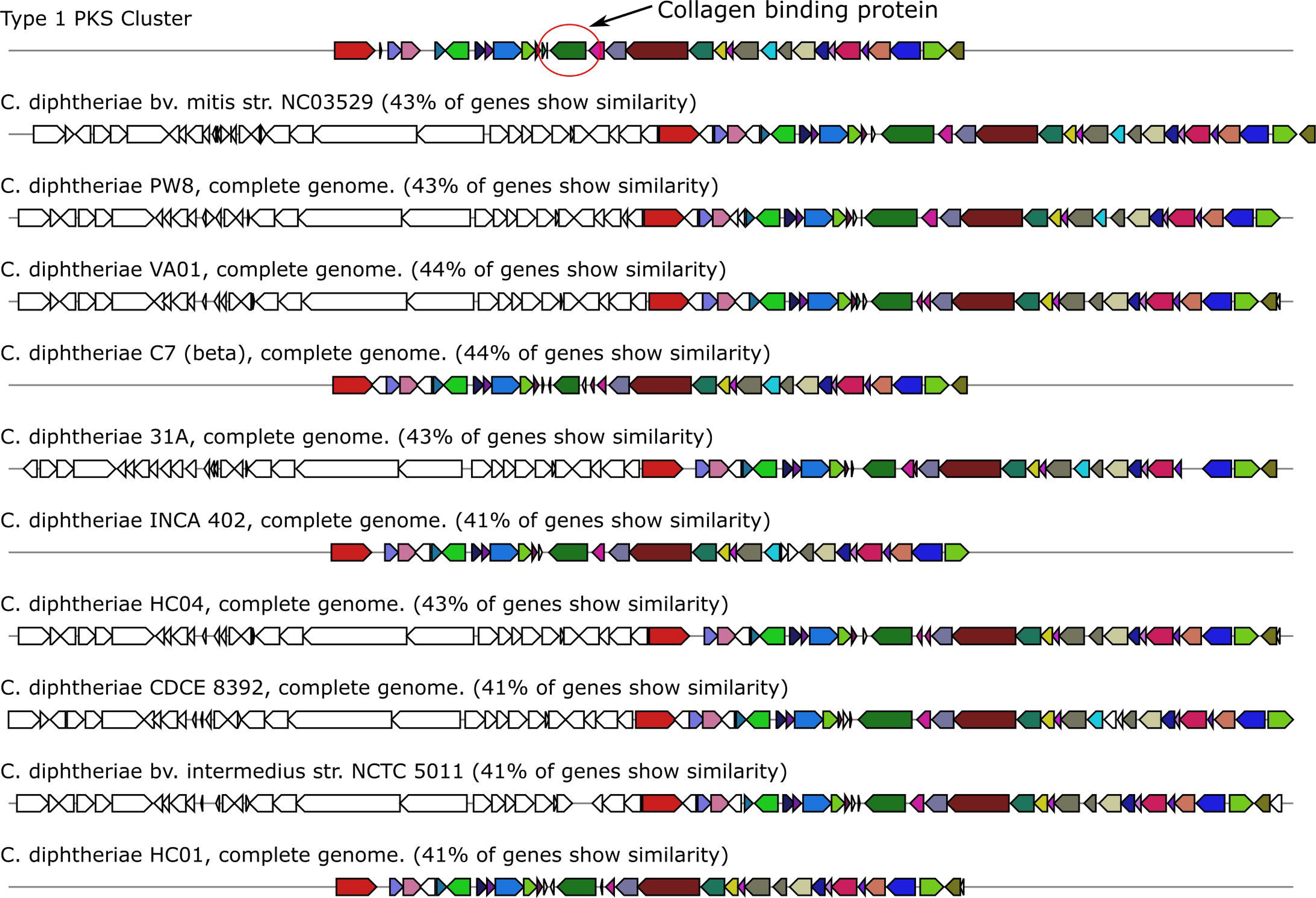
The type 1 PKS cluster of *C. diphtheriae*. The collagen binding protein is indicated by the red circle and corresponds to the same green gene in each cluster. The homology of the cluster is indicated for a selection of strains with the closest homology across the cluster.

Genome-wide comparison of isolates also uncovered concurrent infection in one patient with two genomically distinct strains of *C. diphtheriae*. The first isolate 2767A was toxin gene PCR negative and biotype *gravis* while the second isolate 2767B was toxigenic and biotype *mitis* (Table 1). The isolates were not related to each other according to the analysis employed in this study (Figures 1 and 2).

## Discussion

This is the first report describing the epidemiology of toxigenic and non-toxigenic *C. diphtheriae* in Australia. Apart from the clusters described above, most strains represented very diverse STs reflecting multiple sources of infection including overseas-acquired cases. There is little data on the evolution of *C. diphtheriae*, particularly on current *tox*^-^ strains. We investigated the variability of genomes of toxigenic and non-toxigenic strains of *C. diphtheriae* using a set of isolates collected over the last 12 years in NSW, Australia. While the number of notifiable cases has remained low during this period, the number of isolates referred to our laboratory for diphtheria toxin testing has risen remarkably with 36 of the 47 isolates collected in 2014-2016. This has also been reported by other developed countries and like this study, most isolates are found to be non-toxigenic (18).

This study adds important insights into the evolution of *C. diphtheriae*, particularly on current *tox*^-^ strains in countries with high uptake of diphtheria toxoid vaccines. As immunisation is achieved using vaccine containing diphtheria toxoid, it does not confer immunity to non-toxigenic strains. Infection with non-toxigenic *C. diphtheriae* can still result in respiratory, cutaneous or invasive infections (1, 19) with the cutaneous route speculated to be the more efficient in transmission (20). Proposed risk factors for infection with non-toxigenic strains of *C. diphtheriae* are intravenous drug use, alcoholism and the presence of bone or joint infections, however this is not always the case (21). Furthermore, our findings highlight a potential role of pilins, adhesion factors and iron utilization in infections caused by toxigenic as well as non-toxigenic strains.

The investigation of population structure using traditional MLST profiling, SNP-based mapping of raw reads and pangenome analysis has identified apparent clusters of cases of infection and indicated local acquisition of *C. diphtheriae*. For example, ST32, a ST suspected to have enhanced virulence, was found in four patients. All patients found to have ST32 were from the same region and had a similar age range (16-27). ST32 has been reported previously and the success of this clone is thought to be due to its superior adherence properties (22). The adhesion rate for ST32 was found to be 7.34 ±2.33%, compared to other *tox*^+^ *C. diphtheriae* strains that have an adhesion rate of 0.34 ±0.05% (23). We therefore examined the pilus gene clusters of our ST32 isolates and reconfirmed that the ST32 strains contained the SpaA, SpaD and SpaH pilus gene clusters although the ST32 isolated in our study had poor homology to SpaD from NCTC 13139. The pilus genes clusters are essential for the establishment of infection, particularly in the respiratory tract, and are contained on a pathogenicity island in *C. diphtheriae* (23). They are also known to vary between strains and possibly contribute to the success of certain clones. SpaA type pili are involved in adhesion to pharyngeal epithelial cells while SpaD and SpaH interact with laryngeal and lung epithelium and are highly heterogeneous across strains (5). Loss of *srtA* and/or genes from *spaB* or *spaC* (all from the SpaA pilus gene cluster) equates to loss of adhesion (24). Interestingly, Cluster 1 was the only cluster that contained all three pilus gene clusters and all strains were recovered from patients with respiratory disease.

Antibiotic resistance in *C. diphtheriae* remains relatively uncommon, however, a recent report (25) found that *C. diphtheriae* isolates showed a decreased susceptibility to penicillin and resistance to tetracycline in Rio de Janeiro (25). Multidrug resistant isolates involved in cutaneous and respiratory diphtheria have also been described (26, 27). Further studies have found penicillin resistance to contribute to treatment failure (28). Our findings suggest that antibiotic resistance is uncommon among *C. diphtheriae* in Australia. Only isolates from Cluster 2 appeared to carry the *tetO* gene, encoding a protein which protects the ribosome from the translation inhibition action of tetracycline.

The emerging role of siderophores as important virulence factors of *C. diphtheriae*, deserves special attention. Iron acquisition mechanisms in pathogenic bacteria are known to contribute to survival under “nutritional immunity” a mechanism induced by the host to reduce pathogen cell replication and growth (29). Siderophores are encoded by large non-ribosomal peptide clusters known as NRPS/PKS clusters which confer survival ability in nutrient variable conditions, particularly in establishing and maintaining infection. We demonstrated variability in the siderophore clusters between the STs. Interestingly, some strains contained an additional gene encoding for a collagen binding protein that had high homology among strains isolated from cutaneous or blood infections but low homology (<84%) among strains isolated from respiratory infections. The contribution of collagen binding protein (in combination with an increase in adherence mechanisms) to the success of particular toxin-negative strains warrants further study.

In conclusion, the genomic diversity of toxigenic and non-toxigenic strains of *C. diphtheriae* in Australia suggests multiple local and overseas sources of infection and colonisation. Core and accessory genomes of *C. diphtheriae* strains colonising different ecological niches have significant differences and virulence mechanisms that modulate their fitness as pathogens. Given the growing numbers of *C. diphtheriae* isolates being identified in diagnostic laboratories it becomes essential to closely monitor non-toxigenic strains of *C. diphtheriae*. The findings of additional phage or virulence factors conferring advantage on *tox*^-^ strains of *C. diphtheriae* can be of public health concern as a vaccinated population would have no immunity to new strains given vaccination immunises against the diphtheria toxoid only.

## Acknowledgments

The authors wish to thank Neisha Jeoffreys and the Pathogen Genomics Team at the Centre for Infectious Diseases and Microbiology. Funding from Population Health and Health Services Research Support Program, Round 4, NSW Health, Australia.

## Transparency Declaration

The authors are aware of no relationships/conditions/circumstances that present a potential conflict of interest.

---

**Supplementary table 1:** The allele and ST designations for *C. diphtheriae* genomes

